# Molecular profiling of bovine primordial germ cell specification and migration onset reveals a conserved program in bilaminar disc embryos

**DOI:** 10.1101/2025.06.04.657951

**Authors:** Carly Guiltinan, Ramon C. Botigelli, Rachel B. Arcanjo, Juliana I. Candelaria, Lawrence F. Lanzon, Justin M. Smith, Gloria Becerra-Cortes, Anna C. Denicol

## Abstract

The goal of this study was to establish the timing and location of bovine primordial germ cell (PGC) specification, as well as the molecular characteristics that differentiate early PGCs from other cells in the embryo. A founder population of PGCs was identified via protein immunolocalization in the posterior epiblast (TBXT+) region at E16, by the unique co-expression of OCT4, SOX17, PRDM1, and TFAP2C with absent SOX2. At E20, PGCs were spread along the allantois and hindgut endoderm, indicative of migration onset. Single-cell RNA-sequencing at E20-22 revealed ten PGCs that were subset using the above markers. Further analysis of this population uncovered conserved and novel markers of the early germline and gave numerous insights into their developmental status, including markers associated with pluripotency maintenance, cell migration, repression of the somatic program, and initiation of epigenetic remodeling. This work establishes the region and transcription factor network of bovine PGC specification at day 16 of embryonic development and the first transcriptomic profile of migratory PGCs in cattle, demonstrating a conserved program with other species that form as a bilaminar embryonic disc leading to gastrulation.

## Introduction

Primordial germ cells (PGCs) are the precursors of eggs and sperm and the cellular link bridging one generation to the next. In mammals, PGCs are induced from a subset of cells in the epiblast, the early embryo’s pluripotent entity, preceding gastrulation. The specification of PGCs is a fascinating process and one of the first cell fate decisions of the embryo, since this population must be protected from the signals that support the differentiation of the somatic germ layers that comprise the forming body. Studies in mammalian models have shown that PGC specification is regulated by a tripartite transcriptional network, with some divergence among species. Namely, in rodents *Prdm1*, *Prdm14*, and *Tfap2c* comprise this network^1–3^, whereas in primates and domestic animals *SOX17* (which is not expressed by rodent PGCs) is the critical specifier that operates upstream of *PRDM1* and *TFAP2C*, while *PRDM14* plays a secondary role^4–9^. In all cases, the gene network of early PGCs is necessary for key processes including maintenance of pluripotency, repression of somatic fate, induction of epigenetic reprogramming, and activation of the germ cell program, which are critical processes to the eventual fertility of an animal^10^. Only a small founder population of PGCs is specified, after which these cells directionally migrate through the embryo to the genital ridges, the precursor structure of the gonads. During this time, PGCs proliferate and undergo epigenetic remodeling regulated by the gene network that also commands specification^11^. Upon arriving at the gonads, PGCs interact with the resident gonadal support cells, which initiates PGC differentiation into sex-specific germ cells^12^.

The mechanisms of PGC specification and development in cattle have been unclear, as there are no studies examining the molecular regulation of germ cell specification in the bovine embryo to date. The earliest embryonic stage at which bovine germ cells have been identified is embryonic day (E) 18 by Wrobel and Süß (1998), with putative early migratory PGCs in the caudal wall of the proximal yolk sac, but this study was limited to histological analysis of tissue non-specific alkaline phosphatase activity^13^. In 2021, Soto and Ross characterized bovine germ cells from days 40-90 of fetal age by immunofluorescence (IF) analysis and around day 50 by single-cell RNA-sequencing (scRNA-seq), offering insight into the transcriptome of germ cells once they have reached the ovary^14^. However, a detailed molecular profile of nascent and migratory bovine PGCs has not been described. The delayed start of implantation of bovine embryos until around E20 provides the opportunity to readily obtain peri-gastrulation stage embryonic discs (EDs) for study of earlier events of germline development in this model. Recent studies have examined PGC specification in species with similar embryology to cattle, including pigs, rabbits, monkeys, and humans^6–8,15–17^, showing conserved features in these processes and highlighting the feasibility of studying these events in the cow. Cattle represent a promising model for insights into human reproductive development – as direct studies of human embryos are extremely limited – due to similarities in the timing of embryonic development and gestation length, as well as sample accessibility because of the advanced assisted reproductive technologies (ARTs) and agricultural uses of cattle. Moreover, better understanding bovine germline development would help in the development of revolutionary ARTs that would allow rapid genetic improvement of livestock, such as in vitro gametogenesis from bovine embryonic stem cells (ESCs)^18–20^.

The goal of this study was to perform the first investigation into the molecular profile of the founder population of bovine PGCs during early embryonic development. First, we identified nascent PGCs in the posterior epiblast of bovine embryos at E16 by immunolocalization of established early germline proteins, showing conservation in the specification network with other species that form as an ED. In addition, we confirmed that bovine PGCs were migrating by E20, by immunolocalization of germ cells in the forming allantois and hindgut endoderm. Finally, we analyzed the whole transcriptome of early bovine germ cells (E20-22) by scRNA-seq. Collectively, these studies have provided the most detailed identity to date of nascent and early migratory bovine PGCs.

## Materials & Methods

All reagents and consumables were purchased from Thermo Fisher Scientific unless otherwise specified. Procedures using animals were approved under UC Davis IACUC protocol #22836.

### Embryo production & collection

Cows were housed at the UC Davis feedlot and provided ad libitum water and feed. An established protocol was used to synchronize ovulation^21^. Pregnancies were established and embryos were harvested at day 16 and 20-22 of development. For the E16 collection, bovine embryos were produced in vitro using commercial medium (IVF Biosciences, Falmouth, UK) following manufacturer protocols. Embryos were cultured for 7 days after fertilization, at which time grade 1 blastocysts^22^ were selected and loaded into straws for embryo transfer. Embryos were transferred into synchronized recipients (6 embryos/cow) into the uterine horn ipsilateral with the ovary that had a corpus luteum. Based on previous experiments^23,24^ and since we recovered the embryos within 9 days of transfer (i.e., several days before attachment begins), we do not expect that this number interfered with normal development. Bovine conceptuses were collected from the uterus at E16 by flushing the uterine contents with saline through a modified 20G French Foley catheter and transported to the laboratory on ice within ten minutes. For the E20-22 collection, pregnancies were produced via artificial insemination using X-sorted semen following estrus synchronization. After slaughter of recipient cows, reproductive tracts were transported to the lab within 2 hours. The uterus was dissected out and uterine horns were cut distally with sterile instruments to open the lumen, which was gently flushed with cold phosphate buffered saline with 0.2% polyvinylpyrrolidone (PBS+PVP) using a Pasteur pipette. Once the embryo was identified, it was moved with the Pasteur pipette and/or forceps into a Petri dish with a small amount of cold PBS+PVP. Under a stereomicroscope, conceptuses were examined for the presence of an embryonic disc (E16) or embryo proper (E20-22), which were dissected out and processed for IF or scRNA-seq, while a fragment of trophectoderm from each embryo was collected for sexing.

### Sex determination of bovine embryos

Sex of bovine embryos was determined by polymerase chain reaction of the DEAD box helicase 3 gene (*DDX3*), as previously described^25^. Bordering trophectoderm from each embryo was snap frozen in liquid nitrogen (LN2) and stored at -80 °C. Extraction of DNA was performed using the DNeasy Blood and Tissue Kit (69504, Qiagen) following the manufacturer’s protocol, and quantified using a NanoDrop 2000C Spectrophotometer. Amplification of DNA was performed using GoTaq HotStart Polymerase (M5001, Promega) according to instructions, and amplicons were visualized by agarose gel electrophoresis stained with Ethidium Bromide. Genomic DNA from adult bovine testes and ovaries were used as controls. Discrimination between the X and Y chromosomes was determined based on sex-specific differences in amplicon size (208 bp for *DDX3X* and 184 bp for *DDX3Y*)^26^. Primer sequences used to amplify the *DDX3* gene are AGGAAGCCAGGAAAGTAA (forward) and CATCCACGTTCTAAGTCTC (reverse).

### Immunolocalization of bovine primordial germ cells

Bovine embryos (ED with bordering trophectoderm at E16 or embryo proper with immediately surrounding extraembryonic structures at E20) were fixed at 4 °C in 2.5% paraformaldehyde (PFA) overnight and washed three times and then stored in PBS+PVP at 4 °C. In preparation for cryosectioning, fixed samples were transferred into 30% sucrose at 4 °C until they sank to the bottom of the tube. Each sample was positioned for sagittal sectioning and embedded in Optimal Cooling Temperature (OCT) gel, following manufacturer instructions. Cryoblocks were stored at -80 °C until sectioning. Tissue sections with 8 µm thickness were prepared on a cryostat and mounted on charged Superfrost Plus microscope slides at -20 °C. Slides were stored at -80 °C in an airtight container until staining.

On the day of staining, cryosections were brought to room temperature, washed two times with Tris-buffered saline (TBS) and one time with distilled water to remove OCT. Antigen retrieval was performed with Tris-based antigen unmasking solution (H-3301, Vector Laboratories, pH 9.0) for 20 min at 93 °C in a Coplin jar in a steamer. After cooling down, sections were washed with 0.025% Triton X-100 in TBS (TBS-T), permeabilized for 10 min with 0.5% Triton X-100 in TBS, and washed with TBS-T. Sections were incubated with blocking buffer (10% normal donkey serum, 1% w/v bovine serum albumin (BSA), 0.3 M glycine in TBS-T) for 1 h at room temperature. Primary antibodies diluted in antibody buffer (1% BSA in TBS-T) were incubated overnight at 4 °C. After being washed with TBS-T, sections were incubated with secondary antibodies and Hoechst 33342 in antibody buffer for 1 h at room temperature in the dark. Slides were washed two times with TBS-T and once in TBS. Finally, sections were mounted with Prolong Gold Antifade (Invitrogen, P36930) under a coverslip, cured overnight in the dark, and sealed with transparent nail polish. Antibody incubations were performed in humidified chambers, and washes were done in Coplin jars three times for 5-10 mi each, unless otherwise specified. Sealed slides were stored at -20 °C. Antibodies and dilutions used for immunolocalization are listed in Table 1.

**Table 1.** Antibodies used for protein immunolocalization analysis of bovine embryos. All antibodies were reconstituted according to manufacturer protocols, when applicable.

| **Primary antibodies** | **Manufacturer** | **Catalog number** | **Dilution** |
| --- | --- | --- | --- |
| Goat anti-OCT3/4 | R&D Systems | AF1759 | 1:100 |
| Goat anti-SOX17 | R&D Systems | AF1924 | 1:250 |
| Goat anti-TBXT | R&D Systems | AF2085 | 1:300 |
| Goat anti-TFAP2C | R&D Systems | AF5059 | 1:500 |
| Rabbit anti-SOX2 | BioGenex | NU833-5UC | 1:300 |
| Rabbit anti-Ki67 | Abcam | Ab15580 | 1:500 |
| Rat anti-PRDM1 | Invitrogen | 14596380 | 1:50 |
| **Secondary antibodies** | **Manufacturer** | **Catalog number** | **Dilution** |
| Donkey anti-goat AlexaFluor 568 | Invitrogen | A11057 | 1:500 |
| Donkey anti-rabbit AlexaFluor 488 | Invitrogen | A21206 | 1:500 |
| Donkey anti-rat AlexaFluor 488 | Invitrogen | A21208 | 1:500 |
| **Nuclear dye** | **Manufacturer** | **Catalog number** | **Dilution** |
| Hoechst 33342 | BD Biosciences | BDB561908 | 1:1000 |

**Table 2.** Sexing results from bovine embryos. Genetic sex of embryos analyzed for germ cells, determined by PCR of genomic DNA for *DDX3* homologs. These results only include embryos in which the epiblast portion was intact and appeared developmentally normal, and that proper sectioning allowed the entire span of the epiblast to be analyzed for presence of germ cell markers. IF = immunofluorescent analysis; scRNA-seq = single-cell RNA-sequencing analysis. Age is in embryonic days.

| **Age-ID** | **Sex** | **Analysis** | **PGCs identified?** |
| --- | --- | --- | --- |
| 16-1 | Undetermined | IF | No definitive |
| 16-2 | Male | IF | Yes |
| 16-3 | Female | IF | No |
| 16-4 | Male | IF | Yes |
| 16-5 | Male | IF | Yes |
| 20-1 | Female | IF | Yes |
| 20-2 | Female | scRNA-seq | Yes |
| 22-1 | Female | scRNA-seq | Yes |

Immunofluorescence images were acquired on a confocal microscope equipped with a high content imaging system (ImageXpressMicro, Molecular Devices, San Jose, CA) using a 20X objective with water immersion and 10% tiled overlap. Image processing was performed with the Meta ImageXpress software (v6.7.2.290, Molecular Devices).

### Sample preparation for single-cell RNA-sequencing

In preparation for scRNA-seq analysis, embryos were collected with as little trophectoderm as possible. The entire E20 embryo was collected to increase the number of cells available for sequencing. We removed the anterior portion of the E22 embryo (forming heart and up), in order to better enrich the sample for PGCs. Embryos were slow frozen in cryovials with 90% fetal bovine serum (FBS) and 10% dimethyl sulfoxide using a programmable slow freezer. Slow freezer settings were as follows: first samples were equilibrated for 20 min at 20 °C, then seeded by touching the liquid-air interface with cotton soaked in LN2, held at -7 °C for 10 min, and then cooled to -38 °C at a rate of -0.3 °C per min. Frozen samples were stored in LN2 until being processed for sequencing.

On the day of scRNA-seq submission, slow-frozen embryos were thawed in a 37 °C water bath and immediately washed three times in room temperature HBSS with calcium and magnesium (HBSS^+/+^). Samples were then individually transferred into a 1.5 mL tube with 1 mL of 37 °C cell dissociation medium comprised of HBSS^+/+^ with 1 mg/mL collagenase type IV (LS004188, Worthington Biochemical Corporation) and 5 U/mL DNase I (12185010, Invitrogen) and finely minced using sterile scissors. The resulting solution was transferred into a 15 mL tube with an additional 1 mL of dissociation buffer, pipette mixed 20x using a P1000 with wide bore tip, and incubated for 5 min at 37 °C in a cell shaker at 150 rpm. The pipette mixing-incubation cycle was repeated two times for a total of 15 minutes of dissociation. Then, 10% FBS was added and the solution was filtered through 70 µm followed by 35 µm strainers into a new tube. The filtered solution was centrifuged at 300 xg for 7 min at 8 °C with low brake. Supernatant was removed and the pellet was resuspended in HBSS without calcium and magnesium with 0.4% bovine serum albumin (BSA; 97061416, Avantor). All tubes used throughout the dissociation were pre-coated with 1% BSA to prevent loss of cells on the walls. Cell suspensions (n=2) were submitted to the UC Davis DNA Technologies and Expression Analysis Core facility for counting and viability assessment (Luna-FL, Logos Biosystems) and preparation for scRNA-seq analysis.

### Single-cell RNA-sequencing & bioinformatic analysis

Cell suspensions were processed for library preparation using the 10X Genomics Chromium X microfluidics technology (v3.1) with a targeted recovery of 10,000 cells per sample and sequenced as pair-end 150 bp reads (40,000 reads per cell) using the AVITI platform (both samples on one lane). Alignment of sequencing reads to the bovine genome (NCBI ARS-UCD2.0 with manual *SOX17* annotation from Ensembl ARS-UCD1.2) and initial analyses were performed using the CellRanger pipeline (v8.0.0, 10X Genomics), followed by quality control and secondary analyses using the Seurat package (v5.1.0) of R. Samples were filtered based on nFeatures between 250 and 7,500, nCount <50,000, and <5% mitochondrial genes. Since this cell population was largely comprised of differentiating cells, we regressed out only the difference between the G2M and S phase scores. Data was normalized with the LogNormalize method and a scale factor of 10,000. After identifying variable features by variance-stabilizing transformation and scaling the data, dimensionality reduction was performed by principal component analysis. Integrative analysis was performed using the Harmony integration method. Based on determination of fit by ElbowPlot, we clustered cell populations using the FindNeighbors (dimensions = 1:23) and FindClusters (resolution = 0.2) commands, and then visualized the data using the uniform manifold approximation and projection (UMAP) method (dimensions = 1:23). Cluster identity was assigned using the 10 most differentially expressed genes (FindAllMarkers) and conserved marker genes.

Identification of PGCs was performed by subsetting cells with positive expression (>0.1) of the early germline markers *OCT4*, *SOX17*, *PRDM1*, and *TFAP2C*. Sample age and PGC identity were added as columns to the metadata to allow the integrated object to be split by these factors for comparative analyses. Expression of genes of interest among cell clusters or identities was visualized with FeaturePlot, VlnPlot, and DotPlot. Comparison of genes that encode for cell surface markers was performed between PGCs and soma (i.e., non-PGC lineages of the embryo proper). Direct comparisons of expression of epigenetic and proliferation marks, including cell cycle, between the germline and somatic cells was performed after subsetting the cluster that contained PGCs.

## Results

### Bovine primordial germ cells are specified around day 16 of embryonic development

Bovine conceptuses collected at E16 were filamentous structures with EDs that spanned the pre- to early-primitive streak stages (Fig. 1A). Asynchronous embryonic growth has been widely observed for species including cattle^23,24^, so we attributed the span of stages captured within the same timepoint to normal differences in rate of early development. Homogeneous expression of the pluripotency protein OCT4 throughout the epiblast indicated that the entire structure was still pluripotent at this stage, but axial patterning had occurred, with SOX2 localized to the anterior epiblast and TBXT in the posterior epiblast of all embryos analyzed (Fig. 1B-C). From this pattern of protein expression, we inferred that we had captured bovine embryos just before or at the onset of gastrulation: the period at which we hypothesized PGC specification would occur in cattle based on studies of these events in other mammalian species^6–8,27^.

**Figure 1.**
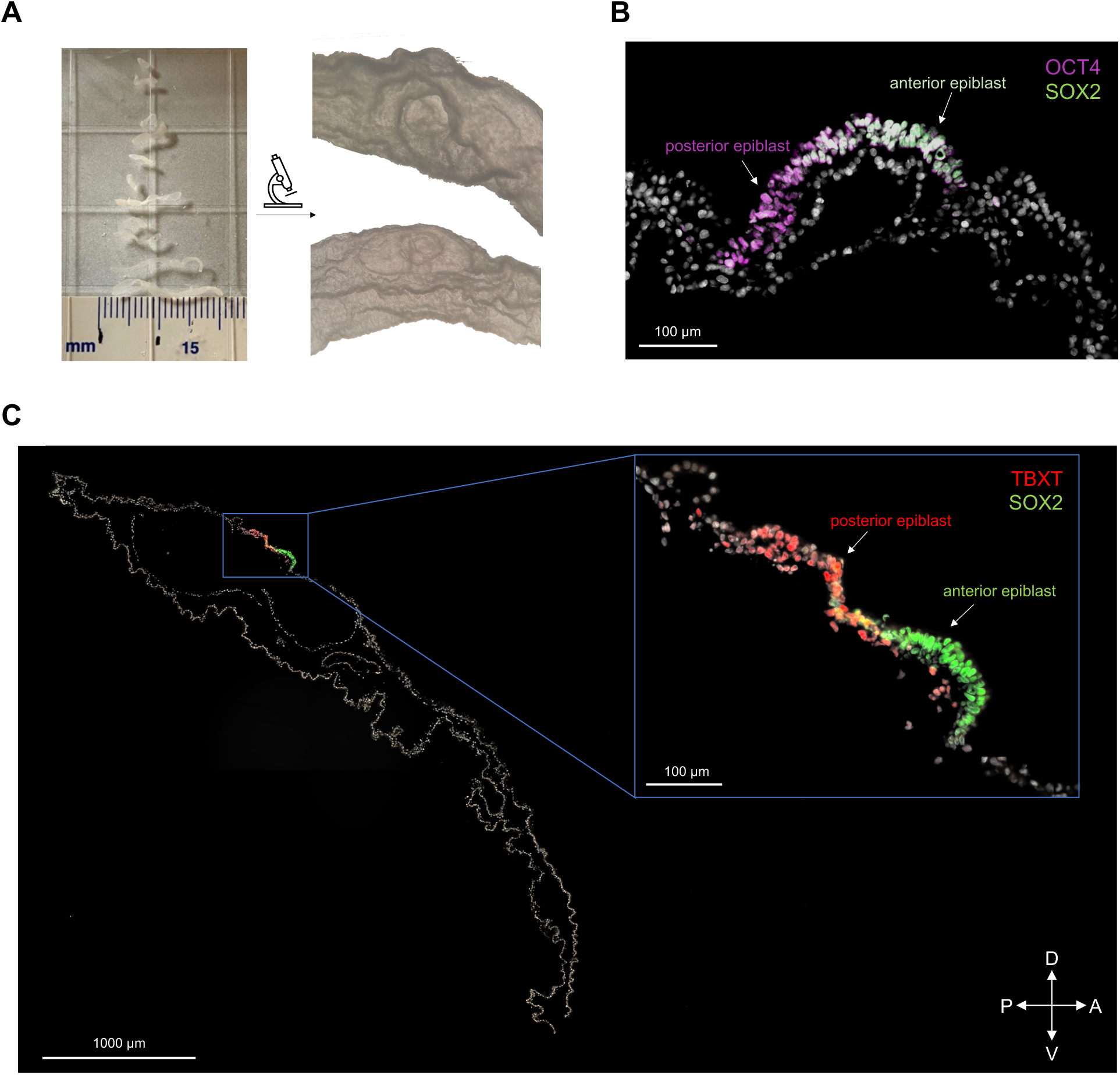
Developmental status of bovine embryos at embryonic day 16. **(A)** Images taken during collection, including embryos or fragments after flushing, and embryonic discs and bordering trophectoderm captured under the microscope. **(B)** Sagittal section of a pre-primitive streak embryo co-stained for OCT4 (magenta)/SOX2 (green). Region with only OCT4 expression is the posterior epiblast, whereas region with OCT4 and SOX2 co-expression is the anterior epiblast. Cells tagged only by DNA dye (gray) are extraembryonic structures. **(C)** Stained sagittal sections of bovine embryos including co-localization of TBXT (red) and SOX2 (green), with TBXT localized in the posterior epiblast and SOX2 in the anterior epiblast, and OCT4 (magenta) present throughout the epiblast. Bottom right key indicates orientation as posterior (P), anterior (A), dorsal (D), and ventral (V).

Consistent with this hypothesis, we identified PGCs in the posterior epiblast region in three out of five E16 embryos by the unique co-localization of early germline markers (SOX17, PRDM1, TFAP2C; Fig. 2A). Of note, these proteins were expressed in other cell lineages of the embryo, with TFAP2C present in the trophectoderm, SOX17 and PRDM1 in the hypoblast, and TFAP2C and OCT4 in putative pre-amnion cells at the edges of the epiblast (Fig. 2A). Moreover, E16 PGCs expressed OCT4 and TBXT, although expression of these markers in the epiblast was not specific to PGCs at this timepoint (Fig. 2B-C). Interestingly, PGCs tended to express OCT4 more highly than the surrounding cells of the epiblast (Fig. 2C). Embryos without definitive PGCs identified were at less advanced stages of development (Fig. 2D).

**Figure 2.**
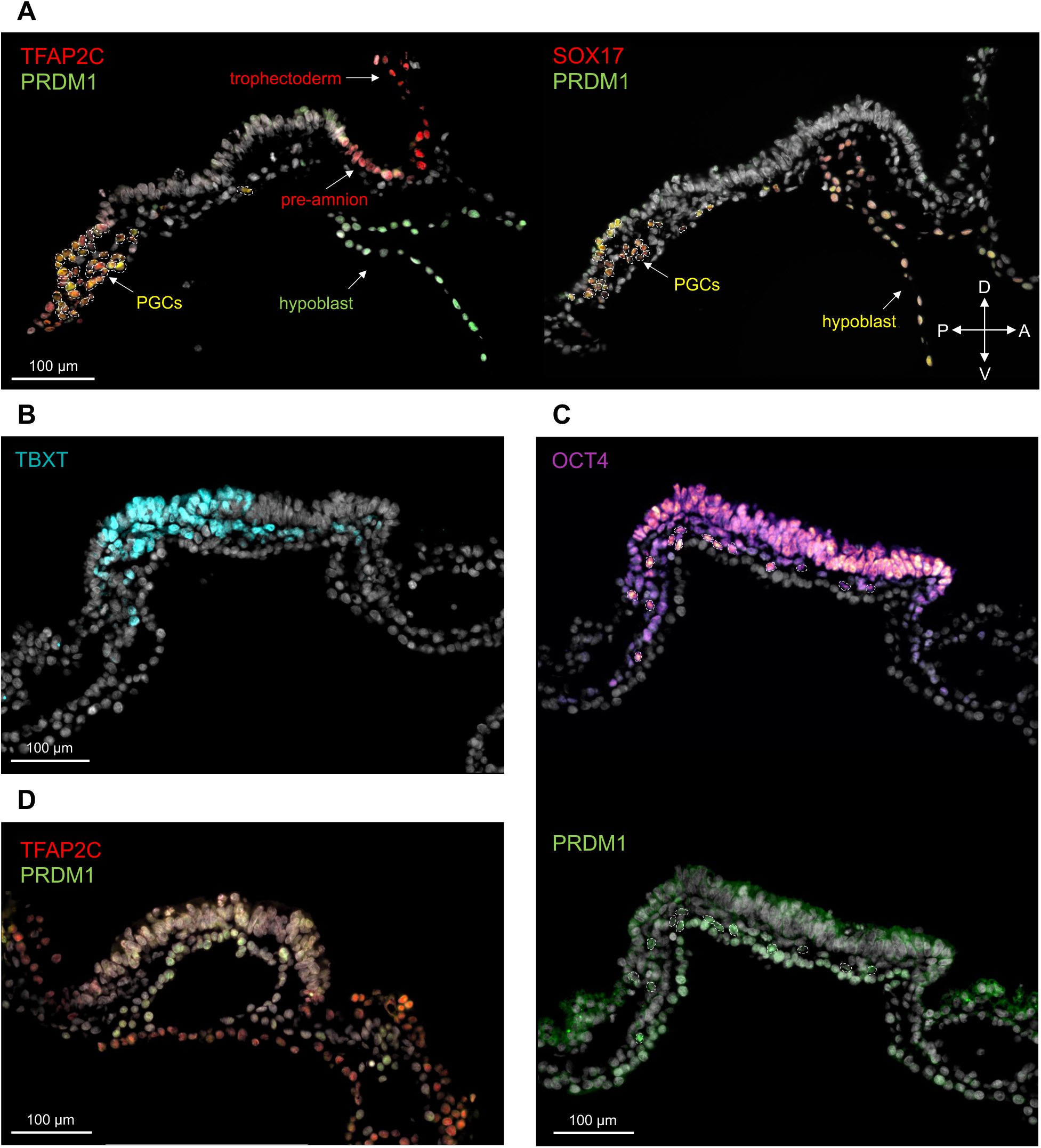
Bovine primordial germ cells emerge around day 16 of bovine embryonic development. **(A)** E16 bovine embryo sections co-stained for early germline proteins TFAP2C (red) and PRDM1 (green) on left, and SOX17 (red) and PRDM1 (green) on right. White dashed circles indicate identified PGCs, with co-expression of these proteins in the epiblast. Bottom right key indicates orientation as posterior (P), anterior (A), dorsal (D), and ventral (V). **(B)** E16 bovine embryo section stained for posterior epiblast marker TBXT (cyan). **(C)** E16 bovine embryo section stained for pluripotency marker OCT4 (hot magenta) on top and PGC marker PRDM1 (green) on bottom. White dotted circles indicate identified PGCs, which co-express OCT4 and PRDM1. **(D)** E16 bovine embryo stained for TFAP2C and PRDM1, with no PGCs identified.

Migration of PGCs initiates with the passive movement of founder cells in the posterior epiblast through the primitive streak^28^. In select sections, we could observe PGCs in the dorsal aspect of the posterior epiblast as well as ventrally, just above where the hypoblast would sit (note: in these sections, the hypoblast was pulled downward; Fig. 2A). We postulate that PGCs in the dorsal region are at their point of origin or moving along the primitive streak, whereas the ventral-most PGCs have already ingressed through along with the developing mesoderm cells, from which population PGCs had been specified, preceded by the cells of the forming endoderm. Collectively, these findings indicate that bovine PGCs are specified in the posterior epiblast by day 16 of bovine embryonic development, and then move through the primitive streak as gastrulation initiates.

### Conserved and diverh2gent features of bovine germline specification with other bilaminar disc-species

Experiments of in vitro germline induction from human ESCs have demonstrated that unlike rodents – in which SOX2 expression is maintained by PGCs and plays roles in early germline development – SOX17 is the key regulator of germline fate while SOX2 is absent^4,5^. Since then, studies in other mammalian species with similar embryology to humans including bilaminar ED formation (e.g., pigs, rabbits, monkeys), have shown that PGCs in these species also lack SOX2 expression and instead rely upon SOX17^6–8^. As anticipated, E16 bovine PGCs were negative for SOX2 and positive for SOX17 protein (Fig. 3A). Along with our other findings, this observation highlights that the tripartite network of specification (i.e., SOX17, PRDM1, and TFAP2C) is conserved among species that form as a bilaminar ED, including cattle.

**Figure 3.**
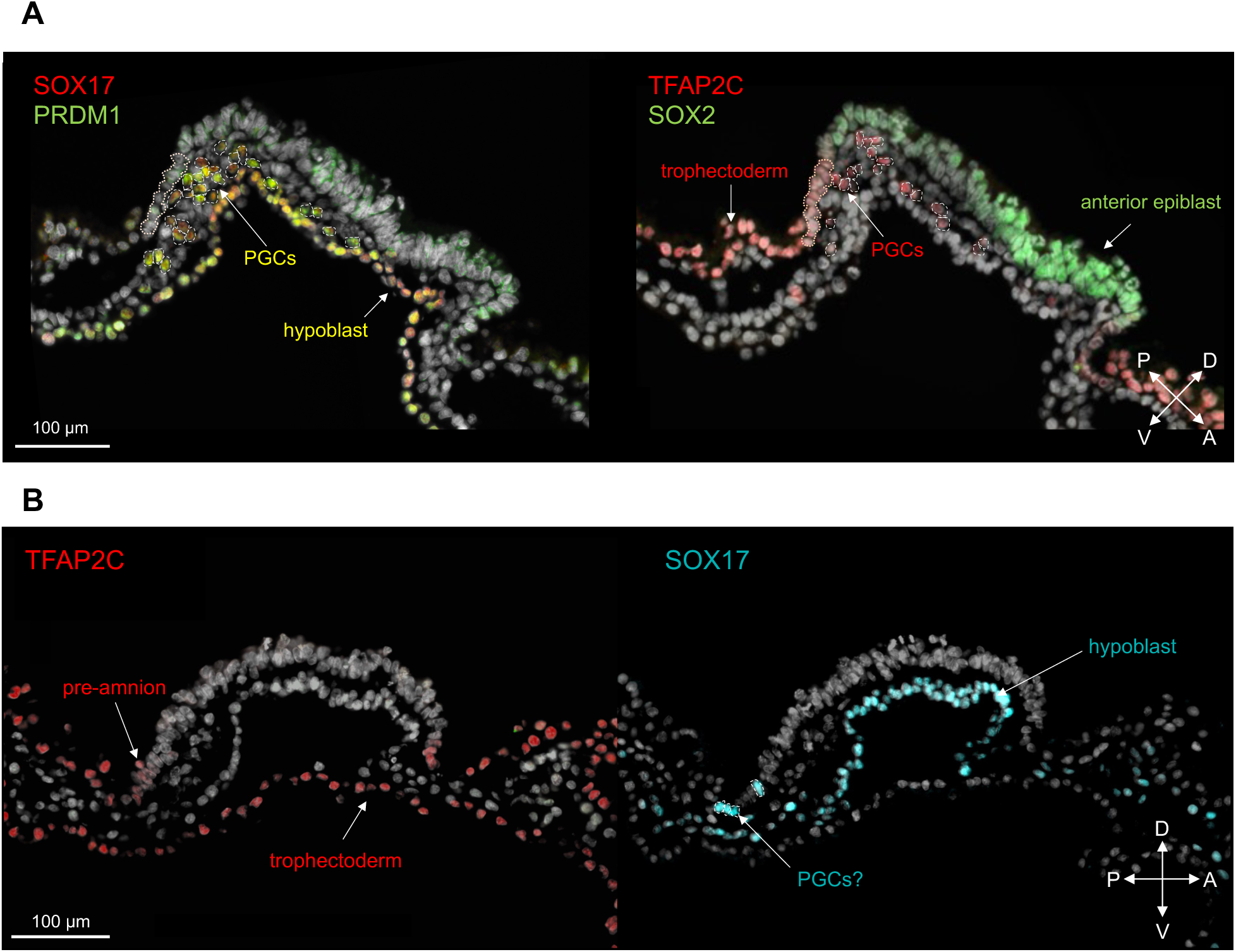
Conserved and divergent features of bovine PGC emergence to other bilaminar disc- species. **(A)** Sagittal sections of E16 bovine embryo stained for SOX17 (red) and PRDM1 (green) on right and TFAP2C (red) and SOX2 (green) on right. White dashed circles indicate identified PGCs, with co-expression of SOX17, PRDM1, TFAP2C, but not SOX2. White dotted region in posterior epiblast shows putative pre-amnion cells. Bottom right key indicates orientation as posterior (P), anterior (A), dorsal (D), and ventral (V). **(B)** Sagittal sections of E16 bovine embryo stained for TFAP2C (red; left) and SOX17 (cyan; right). White dashed circles indicate putative PGCs, which are SOX17+ cells in the pre-amnion region of the posterior epiblast. Bottom right key indicates orientation as posterior (P), anterior (A), dorsal (D), and ventral (V).

Uniquely, studies in cynomolgus monkeys and marmosets have indicated that PGCs are specified from the nascent amnion^6,16^, which has not been implicated in PGC specification of non-primate species. Recent reports have also demonstrated a common progenitor for human amnion and PGC-like cells (PGCLCs) using stem-cell based differentiation approaches^17,29^. The present study shows that bovine PGCs form before advanced development of the amnion, similar to what has been described for pigs and rabbits^7,8^.

We identified putative pre-amnion cells in the embryo sections by co-expression of TFAP2C and OCT4 with negative SOX2 in cells at each end of the epiblast^16,30,31^ (Fig. 2A, 3B). In samples with clear presence of germ cells, we did not observe PGCs among the pre-amnion cells, but rather they were in the posterior epiblast and among the forming mesodermal cells (Fig. 2A). Interestingly, in one of the less developmentally advanced embryos, we identified four SOX17+ cells within the forming pre-amnion cells in the posterior edge of the epiblast (Fig. 3B). This is the first observation of this kind outside of primates, but future cell tracing studies of both lineages would be essential to validate whether the identified SOX17+ cells have a germ cell fate.

### Active migration of bovine primordial germ cells initiates by day 20 of embryonic development

Following passive movement through the primitive streak, PGCs initiate active migration through the hindgut endoderm via the forming allantois^28^. Germ cell migration is facilitated by changes in membrane adhesion to allow movement as single cells and development of pseudopods with chemokine attraction and repulsion to cells along the migratory path to the genital ridges^32–35^. Immunolocalization of bovine embryos at E20 (Fig. 4A) revealed that PGCs were migrating along the forming allantois and hindgut endoderm (Fig. 4B). At E20, PGCs were positive for OCT4, SOX17, PRDM1, and TFAP2C (Fig. 4B-E). Expression of OCT4 in the posterior end of the embryo was exclusive to the germline, highlighting the loss of pluripotency outside of this lineage by day 20, with the exception of the neural ectoderm (SOX2/OCT4-positive) in the anterior region^36^ (Fig. 3B). We also detected PRDM1 in the hindgut endoderm and TFAP2C in the amnion, which had formed by E20, but combined expression of these markers was exclusive to the germline (Fig. 4C).

**Figure 4.**
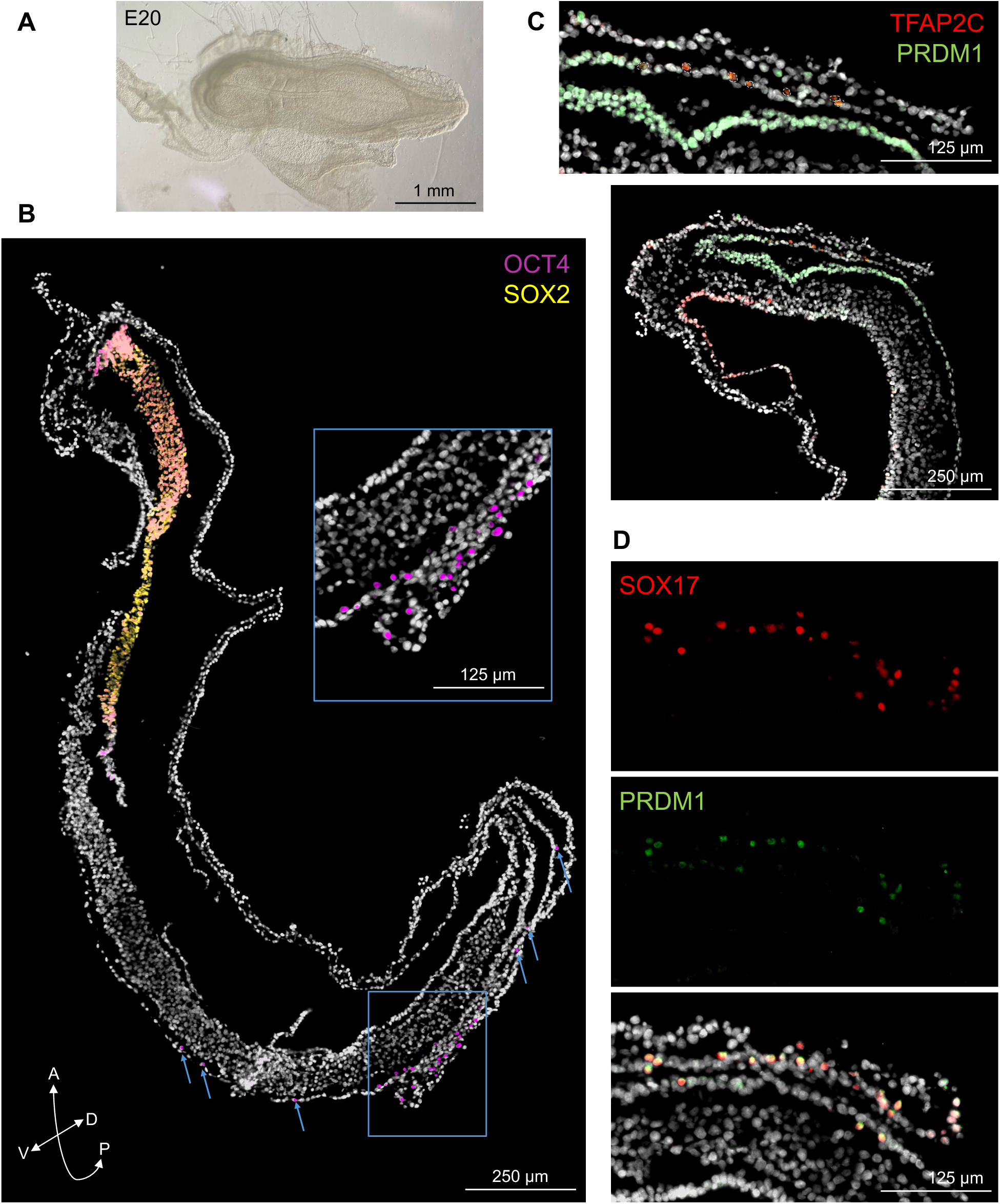
Bovine PGCs initiate migration by E20. **(A)** Stereomicroscope image of E20 bovine embryo. **(B)** Overlay image of stained sagittal section of E20 bovine embryo for OCT4 (magenta) and SOX2 (yellow). Bottom left key indicating orientation as anterior (A), posterior (P), dorsal (D), and ventral (V). Blue inset box and arrows show PGCs (OCT4+). **(C)** Co-localization of germ cell markers TFAP2C (red) and PRDM1 (green) in E20 allantois, including zoomed out image (bottom). **(D)** Co-localization of germ cell markers SOX17 (red) and PRDM1 (green) in E20 allantois, including separate channels and merged image (bottom).

The timing for onset of proliferation of PGCs has been elusive. Studies have shown that some cell division of PGCs takes place during migration, but that the majority occurs upon colonizing the gonads^37,38^. Indeed, active cellular migration exerts high energy, so some have proposed that PGCs alternate between active movement and mitotic division on their journey through the embryo^39,40^. Studies in pigs have found that DNA replication is low in early migratory PGCs, indicating that cells are not yet actively dividing^41^. We immunolocalized Ki67, a marker of cell proliferation, at E20 in order to assess whether early migratory PGCs were mitotically active. Interestingly, while the surrounding cells of the E20 embryo were largely positive to Ki67, 10/13 PGCs were negative (Fig. 5). This suggests that there may be some level of proliferation within the germline shortly after its specification, but that for the most part at E20 PGCs are focused on initiation of cellular migration.

**Figure 5.**
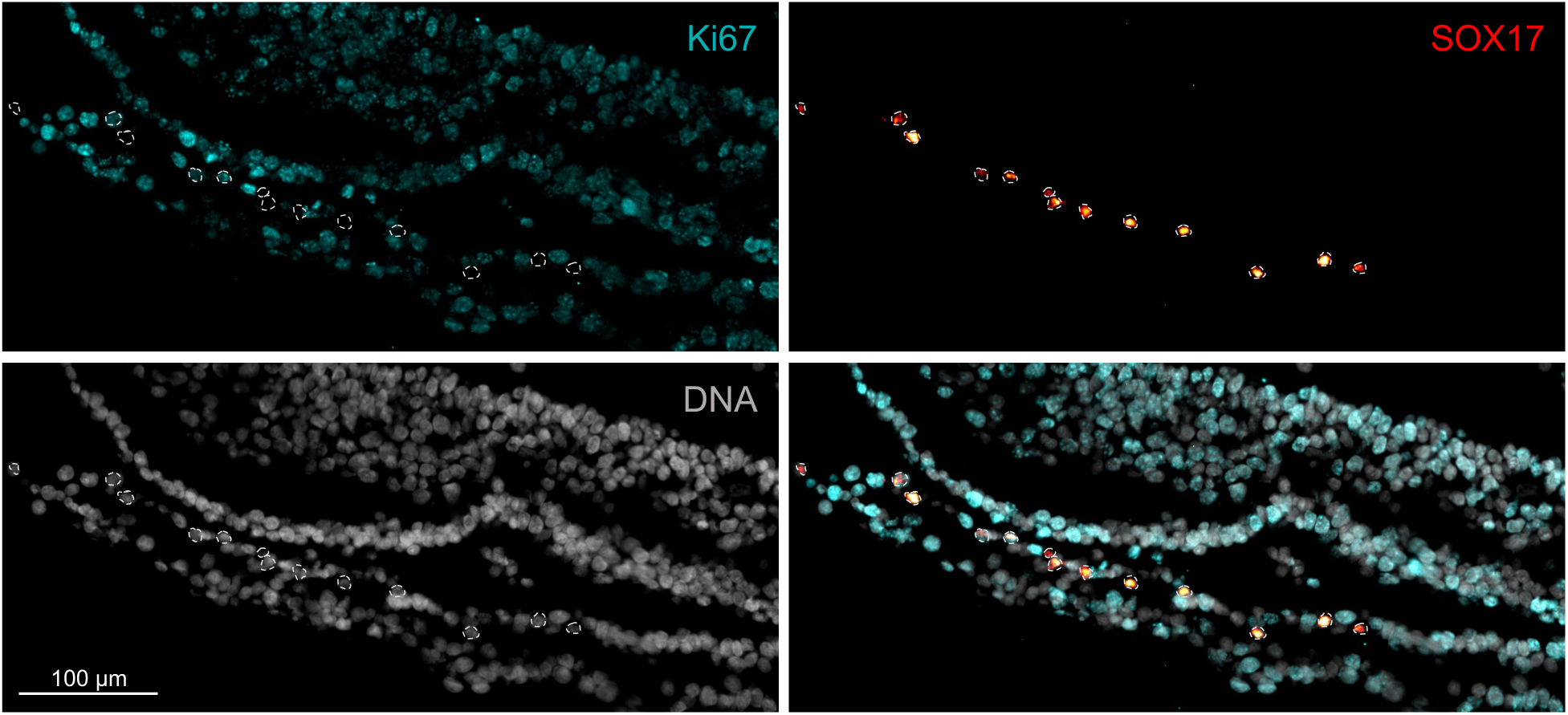
Early migratory bovine PGCs are not highly proliferative. E20 bovine embryo section with co-localization of germ cell marker SOX17 (red hot) and proliferation marker Ki67 (cyan) in the allantois.

### Transcriptome of day 20-22 bovine germ cells revealed by single-cell RNA-sequencing

Next, we analyzed two female bovine embryos at E20-22 at the single cell resolution via scRNA-seq. We saw notable differences in developmental hallmarks between the timepoints, which corresponds with the difference in age (Fig. 6A). Specifically, the E20 embryo had advanced primitive streak development and neural fold formation (2.5 mm in length), while the E22 embryo had completed somitogenesis and body elongation (6.5 mm). In order to enrich for PGCs in the later sample, we removed all structures anterior to the forming heart. Following cell dissociation (56.2-56.6% viability), approximately 16,500 live cells from each sample were submitted for library preparation and sequencing.

**Figure 6.**
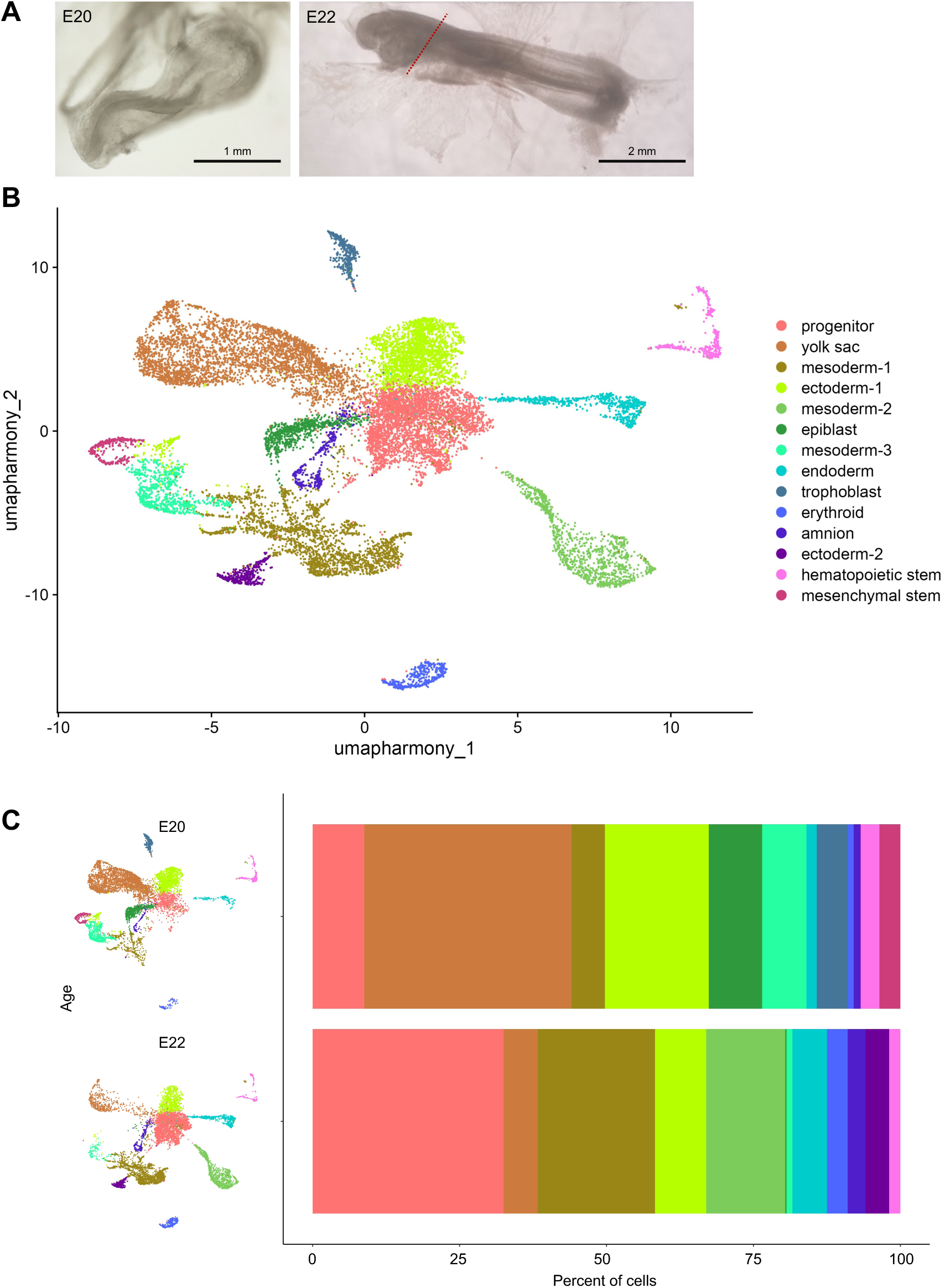
Single-cell RNA-sequencing analysis of bovine embryos at embryonic day 20-22. **(A)** Microscope images of embryos processed for scRNA-seq. Structures anterior of red-dashed line for E22 embryo were dissected prior to sample preparation for submission. **(B)** UMAP from integrated E20-22 samples, with population assignment of each cluster in the legend. Each dot represents a single cell. **(C)** UMAP of E20-22 object split by age (left) with bar graph showing relative proportion of each cluster by age (right). Cluster legend is shown in Fig. 6B.

After quality control and filtering, we identified a total of 17,346 cells (8,265 E20 and 9,081 E22). We found 14 unique cell clusters from the integrated dataset, which represented epiblast-derived and extra-embryonic lineages (Fig. 6B). Specifically, we identified populations derived from the endoderm, mesoderm (3 clusters), ectoderm (2 clusters), pluripotent epiblast, mixed progenitor cells, yolk sac, trophectoderm, amnion, as well as cell lineages likely forming within the yolk sac including primitive erythroblasts, hematopoietic stem cells (HSC), and mesenchymal stem cells (MSC). Top differentially expressed genes used for cluster identification are shown in Figure 7. There was some discrepancy in the cell populations represented in each sample, as expected given the differences in developmental progress (Fig. 6C). Namely, the E22 embryo had evidence of more advanced development and cell differentiation, with derivatives of the three germ layers present and largely absent expression of pluripotency markers. On the other hand, the E20 sample still had a pluripotent entity, consistent with anterior epiblast/forming neural ectoderm based on *SOX2* expression^36^, and comparatively less advanced development of the germ layers, but some representation of the three lineages.

**Figure 7.**
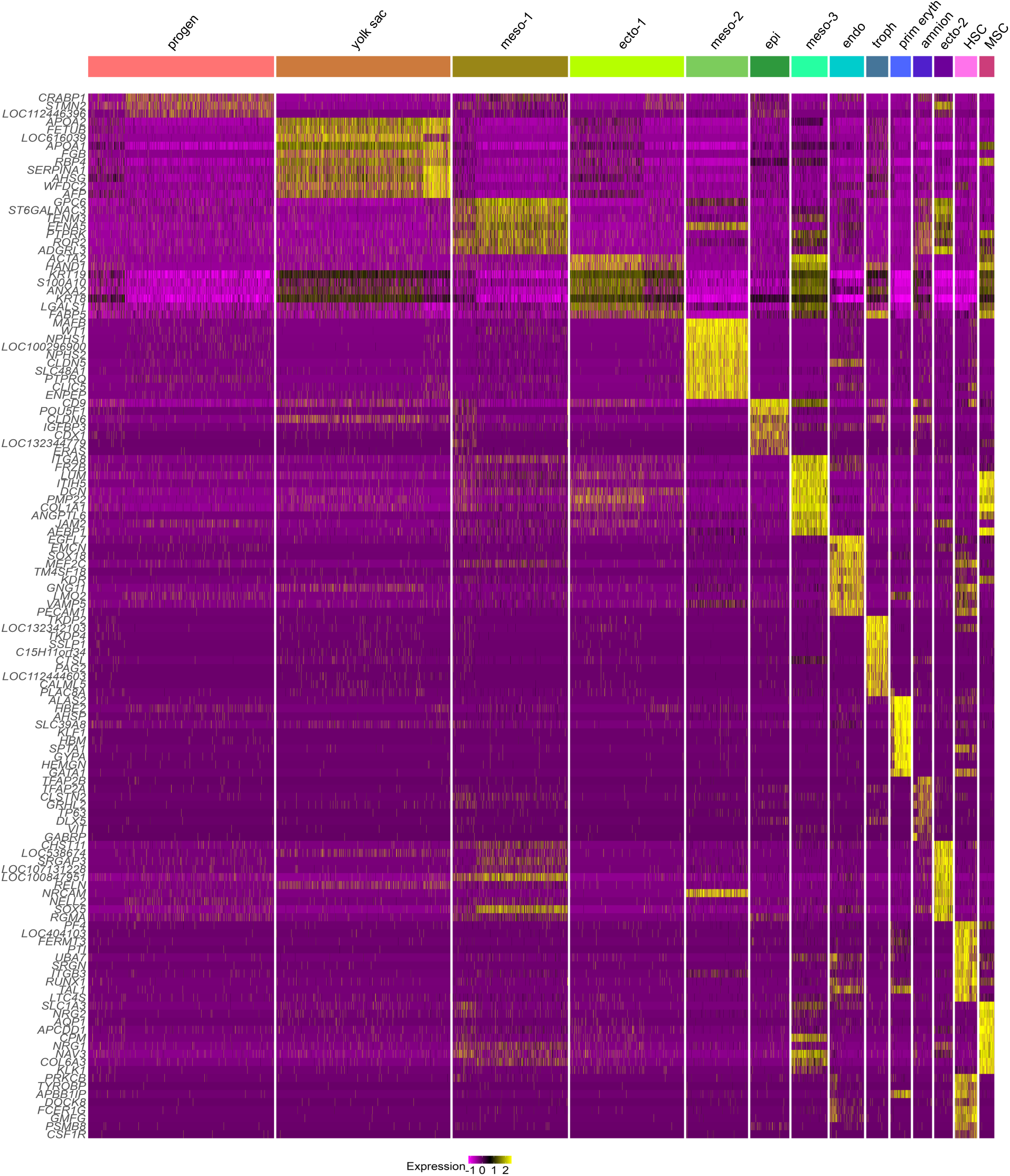
Differential gene expression of cell populations from E20-22 bovine embryos. Heatmap showing top 10 differentially expressed genes with log2(fold change) ≥ 1 for each cluster. Markers were used for assignment of cell type for clusters generated by UMAP analysis.

Considering the small number of PGC founders, we did not detect a cluster comprised purely of germ cells. To identify PGCs within the dataset, we subset cells positive for expression of *OCT4*, *SOX17*, *PRDM1*, and *TFAP2C* (observed at the protein level by IF analysis) from the integrated sample. Of the 17,346 cells sequenced, we captured 10 PGCs (7 at day 20 and 3 at day 22) by these parameters (Fig. 8A). This population expressed markers associated with the germ cell lineage in other mammalian models, including pluripotency (*NANOG*, *KLF4*) and early germline (*NANOS3*, *SOX15*, *DND1*) genes (Fig. 8B). Consistent with our protein staining at day 20, PGCs did not express

**Figure 8.**
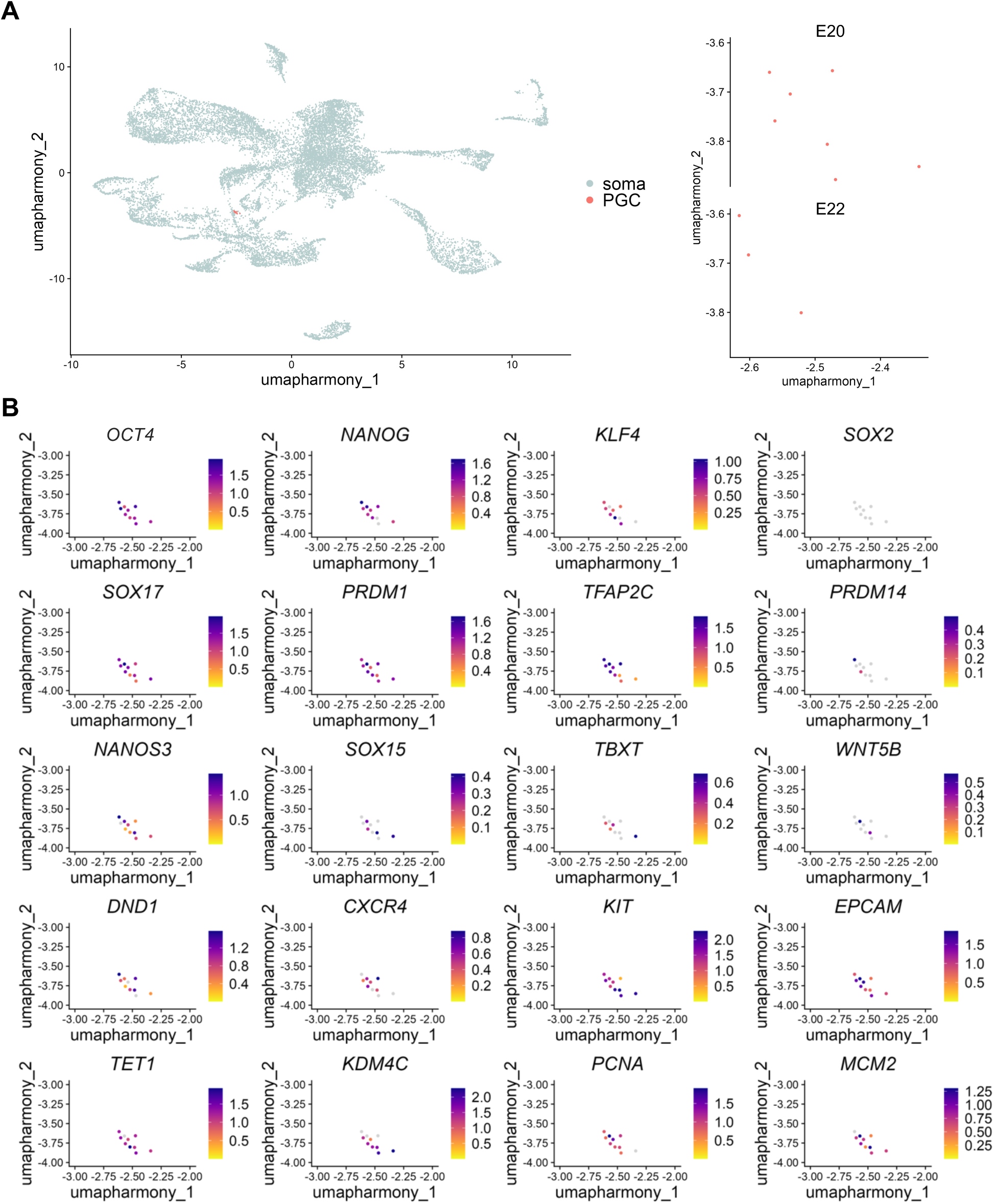
Primordial germ cells identified in E20-22 bovine embryos. **(A)** UMAP data from E20-22 embryos separated as PGCs (*OCT4*/*SOX17*/*PRDM1*/*TFAP2C*+) and soma (remaining cells). Right shows PGCs split by sample age. **(B)** Feature plots with expression of pluripotency (*OCT4*, *NANOG*, *KLF4*, *SOX2*), early germline (*SOX17*, *PRDM1*, *TFAP2C*, *PRDM14*, *NANOS3*, *SOX15*, *DND1*), mesodermal program (*TBXT*, *WNT5B*), migration (*CXCR4*, *KIT*, *EPCAM*), demethylation (*TET1*, *KD4MC*), and proliferation (*PCNA*, *MCM2*) genes in bovine PGCs.

*SOX2*. There was also low/absent expression of *PRDM14*, a major player of PGC specification in the mouse, which has been shown to have secondary roles in the human germline (Fig. 8B). Moreover, we identified genes that encode for surface proteins associated with active cellular migration (*CXCR4, KIT, EPCAM*)^42^. At this point, PGCs had low/absent expression of mesodermal genes (*TBXT*, *WNT5*), associated with repression of somatic fate (Fig. 6B). In addition, we saw evidence for initiation of epigenetic remodeling (*TET1*, *KDM4C)* and cell proliferation (*PCNA*, *MCM2*; Fig. 8B). No age-specific trends in relative expression were observed. As expected, PGCs at this stage did not express genes associated with late migration (*CDH5*, *DMRT1*), germline commitment (*DAZL*, *DDX4*), or meiotic entry (*STRA8*, *SYCP1*; data not shown). Overall, there was some heterogeneity in the presence and expression levels of the markers captured, which likely is reflective both of limitations of the sequencing assay and asynchrony of early germline formation.

### Insights into developmental hallmarks of bovine PGCs at the onset of migration

In order to further establish the developmental status of bovine PGCs around E20, we compared cell cycle and epigenetic markers among PGCs and somatic cells of the same cluster. We found that 7/10 PGCs were in G1, 2/10 in G2M, and 1/10 in S phases, compared to 33.6% in G1, 32.4% in G2M, and 34.0% in S for somatic cells (Fig. 9A). Alongside positive expression of *PCNA* and *MCM2* (Fig. 8B), as well as our prior observation that E20 PGCs were negative for Ki67 protein, this suggests that early germline cells are predominantly preparing for proliferation but not yet actively dividing around E20.

**Figure 9.**
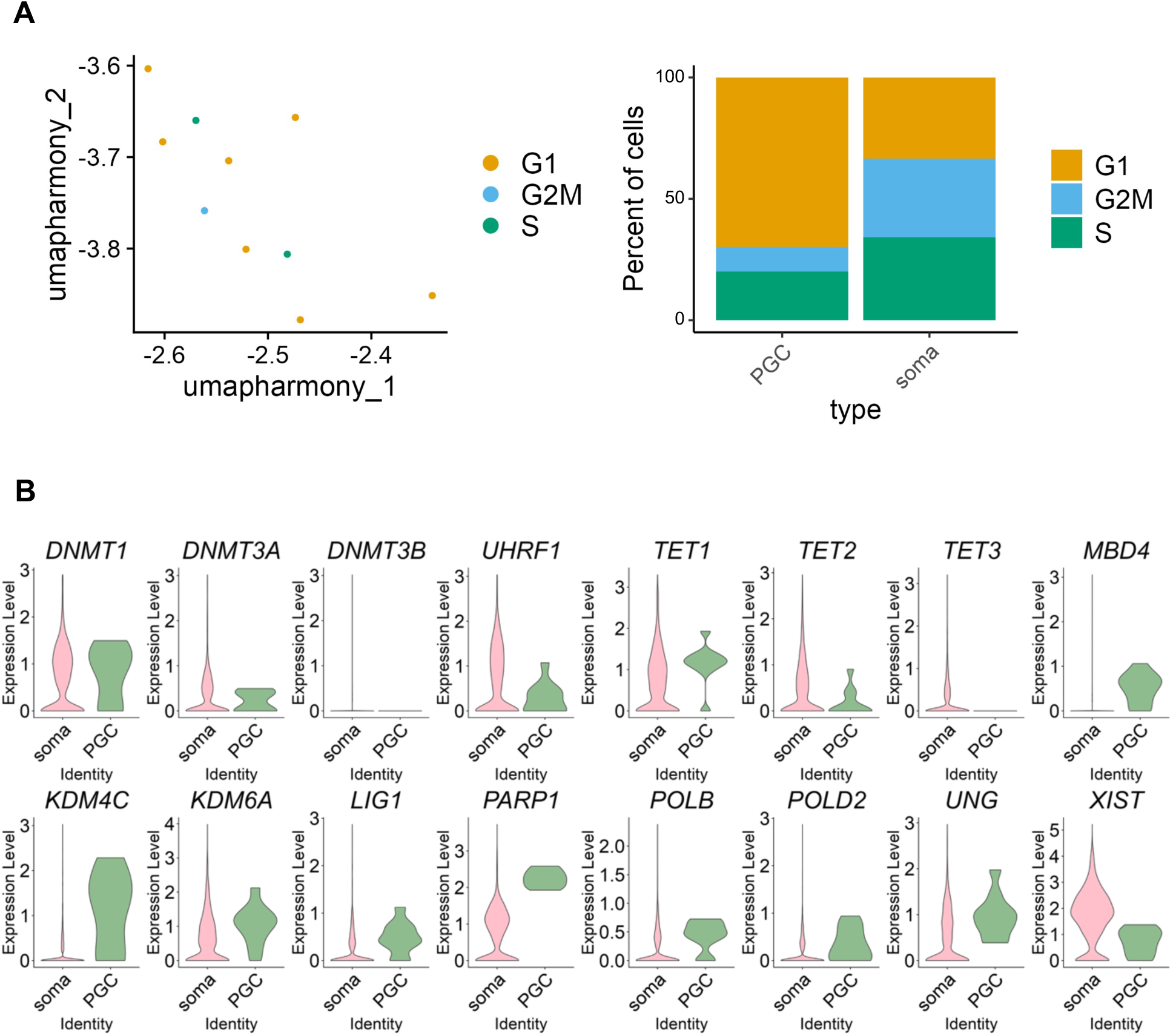
Proliferation and epigenetic status of early bovine primordial germ cells relative to their somatic counterparts. **(A)** UMAP of PGCs split by cell cycle stage (left) and relative proportions compared to somatic cells within the same cluster (right). **(B)** Violin plots showing relative expression of PGC and soma from same cluster for markers for DNA methylation (*DNMT1*, *DNMT3A*, *DNMT3B*, *UHRF1*) and demethylation (*TET1*, *TET2*, *TET3*, *KD4MC*, *KDM6A*), base excision repair (*LIG1*, *PARP1*, *POLB*, *POLD2*, *UNG*), and X-chromosome inactivation (*XIST*).

Following their specification, PGCs undergo extensive epigenetic remodeling, including global DNA demethylation^11,43,44^. This cellular reprogramming is critical for the erasure of parental marks and gives PGCs competency for their continued development into gametes^45^. We assessed the expression of markers associated with DNA methylation (*DNMT1*, *DNMT3A*, *DNMT3B*, *UHRF1*), demethylation (*TET1*, *TET2*, *TET3*, *KDM4C*, *KDM6C*), base excision repair (*MBD4*, *LIG1*, *PARP1*, *POLB*, *POLD2*), and X-chromosome inactivation (*XIST*) in E20 PGCs vs. somatic cells of the same cluster (Fig. 9B). The TET enzymes are involved in catalyzing the conversion of 5mC to 5hmC, which occurs upstream of DNA demethylation and has been shown in the germline shortly after specification^41^. Germline expression of *TET1* and *TET2* suggest that this conversion process has initiated in bovine PGCs by E20. In addition, expression of methylation markers was lower in the germline, while DNA demethylation markers had elevated expression levels (Fig. 9B). We also identified upregulation of several markers of base excision repair – which plays roles during active DNA demethylation – in the E20 germline compared to somatic cells, similar to observations in E14 porcine PGCs^41^. Upon specification, PGCs regain features of naïve pluripotency, including X chromosome reactivation in females. Studies in mice have shown that X reactivation is a gradual process, which starts in nascent PGCs by cessation of XIST expression, a marker of X chromosome inactivation^46^. As well, evidence of X chromosome reactivation has been shown in early-pre-migratory porcine PGCs^41^. Compared to somatic cells, PGCs had lower expression of *XIST* transcript (Fig. 9B), indicating that X reactivation initiates in female bovine PGCs shortly after their formation. Taken together, these findings indicate that epigenetic reprogramming of the bovine germline initiates around the onset of migration.

### Novel and highly specific marker of early mammalian primordial germ cells, RBMXL2

Investigation of top upregulated genes in PGCs vs. somatic cells of the embryo uncovered that the top upregulated gene by fold change (log2FC = 9.73, p_adj = 3.0×10^-238^) was *RBMXL2*. Upon comparing the expression of this transcript in the E20-22 bovine embryo, we found that it was highly specific to the germline (Fig. 10A), with comparable expression levels only in a small number of cells from the epiblast, which were exclusive to the E20 sample. The function of RBMXL2 was only recently elucidated, and studies of this factor have been limited. Ehrmann et al. (2019) first demonstrated that RBMXL2 is a germline-specific RNA-binding protein that functions to repress aberrant splicing patterns during meiosis^47^. It is encoded by a retrogene, *RBMXL2*, which is conserved among mammals^48^. The authors showed that deletion of *RBMXL2* blocks spermatogenesis in the mouse, and that its roles are essential for progression through prophase I in the adult male. Moreover, a recent human case report showed that a frameshift mutation in *RBMXL2* led to cryptic splice poisoning and meiotic arrest during spermatogenesis, leading to infertility^49^. To our knowledge, *RBMXL2* has not been implicated in early germline development during embryogenesis. Interestingly, PGCs expressed both *RBMXL2* (Fig. 10A) and the somatic cell-expressed counterpart, *RBMX*^50^ (Fig. 10B). It is possible that E20 PGCs are transitioning from expression of *RBMX* to *RBMXL2* as they are acquiring the germ cell fate, or that only one of these factors is actively translating protein. Although no markers associated with meiotic entry or progression were detected, as anticipated since this process does not begin until around 75 days of gestation in cows, this observation supports the notion that the germline is already preparing for advanced stages of development shortly after its formation.

**Figure 10.**
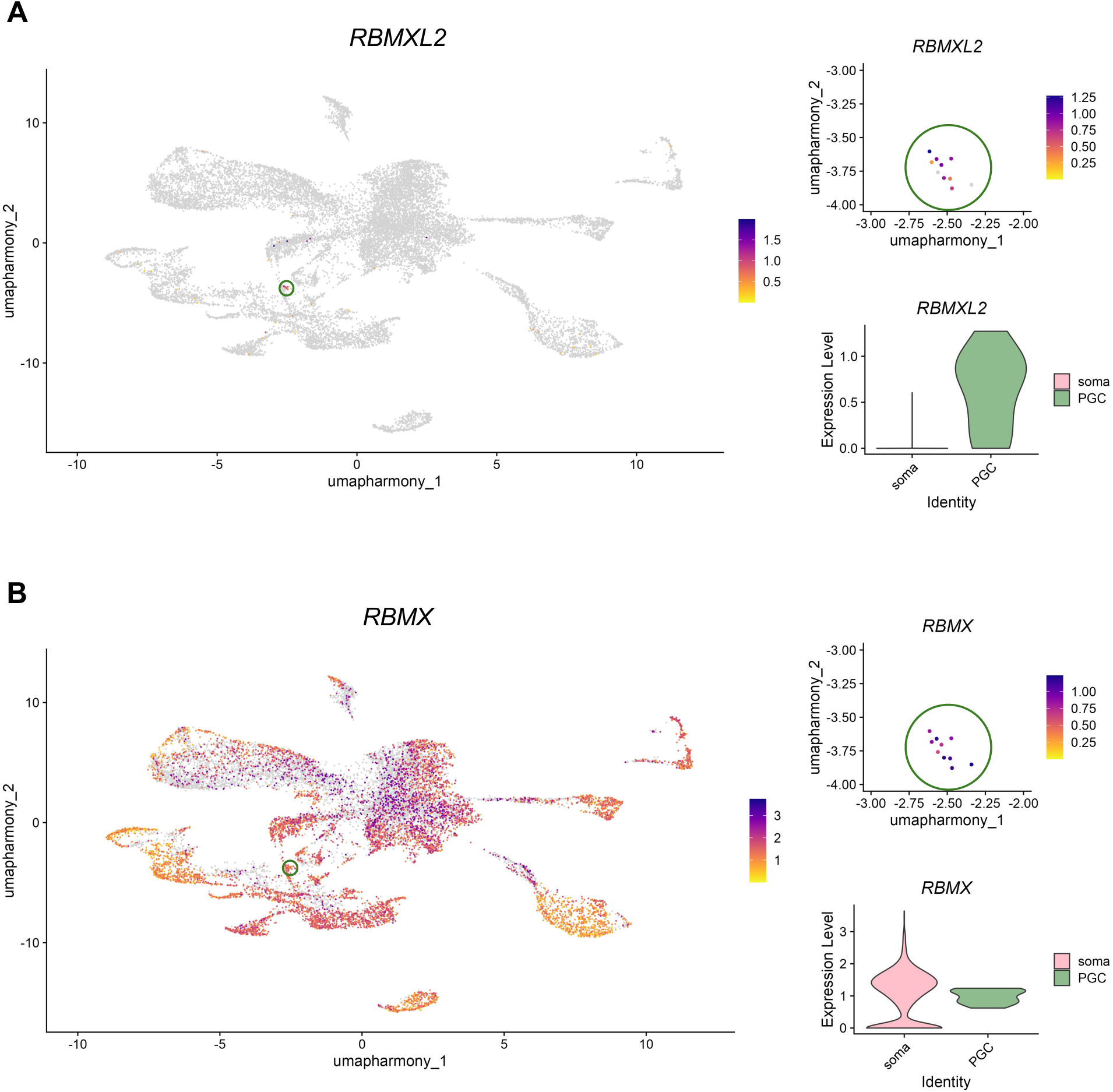
Identification of a novel and highly specific early germline marker, *RBMXL2*. **(A)** Feature plot showing expression of germline RNA binding regulation gene *RBMXL2* with green circle indicating region with PGCs (left). Dot plot and violin plot showing expression of *RBMXL2* within PGC population (right top) and expression in PGCs (green) relative to somatic cells (pink) of the same cluster (right bottom). **(B)** Feature plot showing expression of RNA binding regulation gene *RBMX* with green circle indicating region with PGCs (left). Dot plot and violin plot showing expression of *RBMX* within PGC population (right top) and expression in PGCs (green) relative to somatic cells (pink) of the same cluster (right bottom).

### Candidate cell surface markers of bovine primordial germ cells and in vitro-derived counterparts

Strategies for in vitro gametogenesis from pluripotent stem cells rely on purification of PGCLCs from non-induced cells before re-aggregation or co-culture with somatic cells of the fetal gonad^51,52^. Moreover, efficient isolation of PGCs from bovine embryos would facilitate further studies with these cells, which has yet to be shown. In order to obtain viable cells with high purity, a specific cell surface protein – or more often, a combination – must be determined. We screened a panel of genes that encode for cell membrane proteins, including top upregulated genes identified in bovine PGCs and markers previously used for PGC and PGCLC sorting strategies from other mammalian species^4,5,41,53–56^. We compared the proportion of cells among PGCs and somatic cell populations of the embryo that expressed genes that encode for these proteins (Fig. 11A). Newly identified candidate germline surface markers *CDH4* and *CD83* were expressed in 8 and 9/10 captured bovine PGCs, respectively, while only being expressed in 775 (4.5%) and 239 (1.4%) of the 17,338 somatic cells of the embryo. Although commonly used for sorting strategies in humans, PDPN and ITGA6 were expressed in less than half of E20 bovine PGCs, and were not highly specific, while CD38 was not detected in any early PGCs. Although GPR50 was identified as a specific marker of the E14 porcine germline^41^, we did not identify this marker in bovine PGCs. Shirasawa et al. (2024) isolated bovine ESC-derived PGCLCs by targeting cells that were positive for KIT and negative for CD44 protein^57^. We observed *KIT* in 10/10 and *CD44* in 0/10 PGCs in our dataset, but there were also 1,608 (9.3%) *KIT*+/*CD44*-somatic cells in the embryo dataset, predominantly in populations derived from mesoderm. A migratory protein that participates in chemotaxis during migration, *CXCR4*, has been exhibited as a highly expressed surface marker of human PGCLCs^58,59^, while another recent study identified FGFR3 as a germline protein expressed by PGCs and repressed upon meiotic initiation^60^. Transcripts for *CXCR4* were expressed in 6/10 PGCs, and *FGFR3* in 9/10 PGCs, and represented only 864-904 (5.0- 5.2%) of the other cells captured.

**Figure 11.**
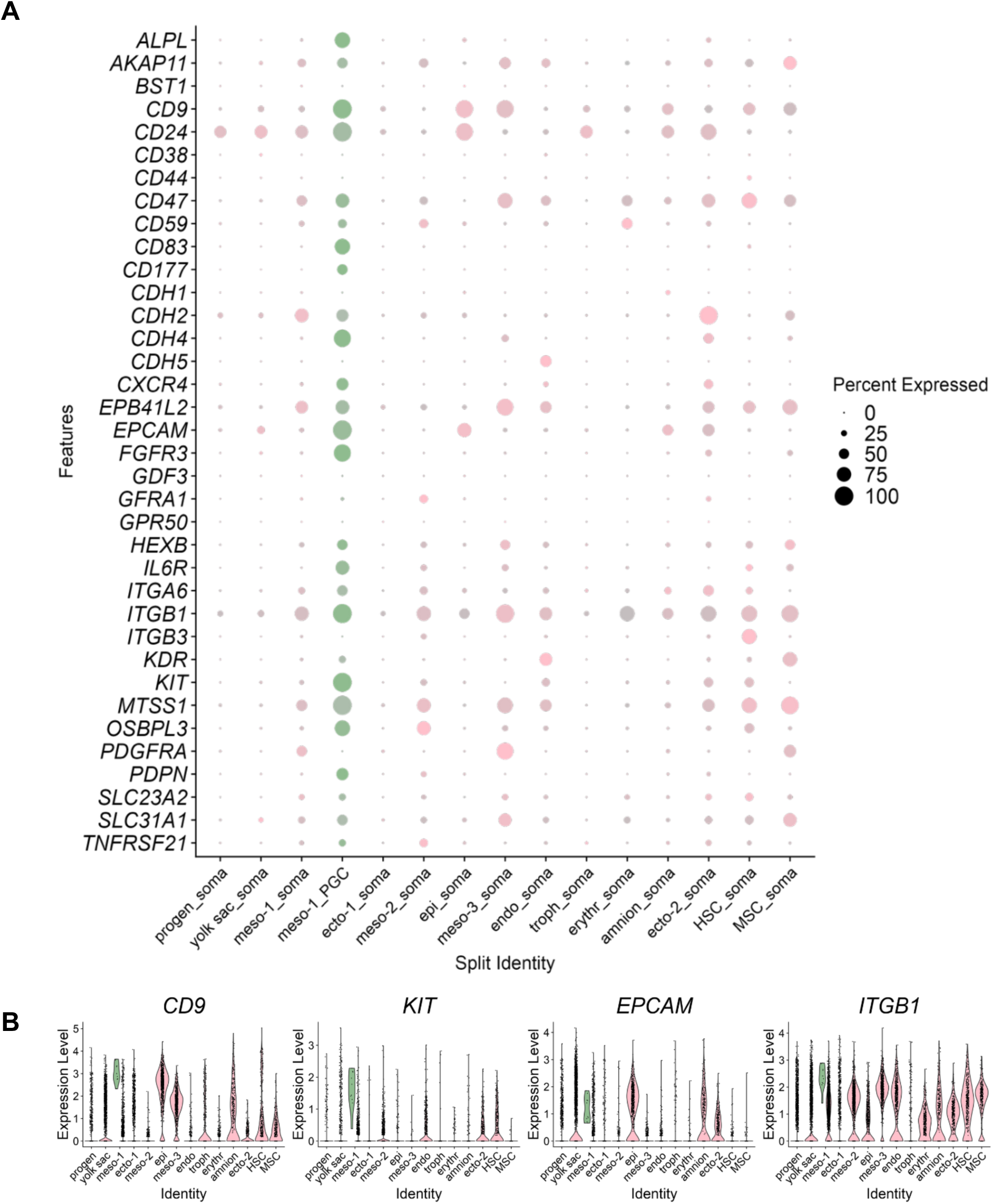
Candidate cell surface markers for primordial germ cell sorting strategies. **(A)** Dot plot showing the proportion of cells in each cluster of E20-22 embryos that express candidate cell surface markers, split by PGC (green) and soma (pink). Circle size indicates percentage of cells within each identity with positive expression; gray scale indicates lower relative expression. **(B)** Violin plots showing expression level of cell surface markers *CD9, KIT*, *EPCAM*, and *ITGB1*, in PGCs vs. somatic lineages of the embryo.

Another potential strategy for cell sorting is targeting proteins by expression level, especially in instances where expression is not highly specific or cells are being sorted from tissues comprised of diverse cell types. We analyzed relative transcript expression level of four surface markers of interest, *CD9*, *KIT*, *EPCAM*, and *ITGB1*, in bovine PGCs vs. other cell types of the embryo (Fig. 11B). Embryonic stem cells are the in vitro-counterpart of the epiblast, so the epiblast cells captured in this analysis would be the closest proxy for the expression profile of bovine ESCs. We observed similar expression levels of *CD9* and *EPCAM* in PGCs and epiblast cells (Fig. 11B). However, relative to non-epiblast lineages, *CD9* and *EPCAM* were markedly higher in the germline, which indicates usefulness for sorting from post-gastrulation embryos. On the other hand, *KIT* and *ITGB1* were upregulated in PGCs relative to the majority of non-germline cells in the embryo, including the epiblast (Fig. 11B). Indeed, bovine ESCs have been exhibited to have low KIT protein expression compared to bovine PGCLCs^57^, corroborating the trend seen in the sequencing data. These findings highlight that cell sorting strategies using the surface proteins encoded by these genes could potentially target cells by expression level in order to purify PGCs with higher fidelity.

## Discussion

Beyond giving novel insights into the timing and regulation of early germline development in the cow, the findings of the present study can be viewed in the context of similar progress with other mammalian species, including rodents, primates, and other domestic animals. Namely, the tripartite transcriptional network that commands specification was conserved among cattle, monkeys, pigs, and rabbits, all of which form as an embryonic disc, and distinct from rodents, which form as an egg cylinder^15^. Moreover, the molecular profile of nascent and early PGCs was similar among ED-forming species, although there were some distinct features, including cell surface marker candidates.

Primates and livestock species have distinct aspects of embryogenesis, including the timing of germline emergence relative to implantation and amnion development^15,61^. We found evidence of PGC-precursors among putative pre-amnion cells (OCT4/TFAP2C+) of the E16 bovine posterior epiblast that did not yet have germ cells present, based on expression of SOX17 – the first-acquired marker of human PGCs^4^. The inferred trajectory of these cells toward the amnion lineage is corroborated by confirmed expression of TFAP2C in the forming bovine amnion at E20. Further supporting this idea, our scRNA-seq analysis shows transcriptomic similarity between the amnion and PGCs, with these populations clustering closely together. While we cannot definitively conclude that the SOX17+ cells have a germline trajectory, this finding warrants further cell tracing studies to test the possibility that the very first PGCs emerge in the pre-amnion region of the posterior epiblast before they move toward the posterior tip of the primitive streak and then ingress through this structure as gastrulation initiates, which is the developmental status captured in our more advanced E16 bovine embryos. We surmise that – similar to the primate^16,17^ – bovine PGCs and amnion cells possibly have a common origin, and then undergo unique developmental trajectories following their specification within the same region, which has yet to be shown in a non-primate species.

Given that the founder population of PGCs is exceptionally small, we did not anticipate capturing a large number of cells by scRNA-seq since we did not highly enrich for them prior to sample submission. Interestingly, we captured a higher number of PGCs (n = 7/8,265) at E20 than we did at E22 (n = 3/9,081). We reason that this is due to lower enrichment of PGCs within the E22 sample, which represents a more advanced and complex embryo. The embryo had grown considerably from E20 to E22, but the same number of cells were submitted for sequencing from both samples. Moreover, our analysis of PGC cell cycle and Ki67 expression show that the cells are unlikely to be rapidly proliferating at this point. Therefore, we attribute the lower number of PGCs at E22 to these cells comprising a lower percentage of the sample. While providing numerous novel insights into bovine PGCs, one important limitation of this study is the low number of cells that were captured by the scRNA-seq analysis. Further studies on a larger number of cells will be crucial for confirming and expanding upon our observations. Additional studies of a larger pool of germ cells could also include integration with other publicly-available datasets, in order to perform side-by-side comparisons of the early germline in various mammalian species and provide new insights into the evolutionary divergence of the germ cell lineage.

This analysis demonstrated for the first time that germline-specific splicing factor *RBMXL2* is expressed in the early germline. We detected co-expression of this factor with *RBMX*, which the somatic cell lineage relies upon for control of splicing and maintenance of genome integrity^50^. Future studies could address whether RBMXL2 is being translated at this stage of development, as this protein has only been identified in adult male germ cells previously, where it functions during meiosis. Although these cells are far from meiotic initiation, *RBMXL2* may function in maintaining genomic stability during earlier events of germline development, such as initiation of epigenetic remodeling, acquisition of the germ cell program, and/or protection against surrounding signaling cascades during migration, which warrants further studies of this gene during early germline establishment. This observation provides novel insights into the potential control of genomic integrity in the early germline.

Increased understanding of the molecular signaling governing germ cell specification in cattle has vast implications for fertility restoration and maximization of reproductive capacity in this species. Advancements provided by these data can be applied to develop efficient approaches to obtain bovine PGCLCs from ESCs, opening a myriad of basic and applied research opportunities, including but not limited to in vitro breeding^19,20^. Importantly, studies of in vitro-generated PGCLCs have indicated that these cells most closely resemble early migratory PGCs^58^. The present study provides a full transcriptional profile of bovine PGCs at this stage of development, providing critical insights into methods to purify and assess cells produced by these approaches, including novel germline markers. Finally, similar studies of developmental biology of bovine embryos in the peri-gastrulation period can answer questions about the other cell fate decisions, which can be applicable to fields of study including in vitro differentiation of bovine ESCs toward somatic cell fates for other studies and applications.

## Conclusions

Collectively, the present studies demonstrate that bovine PGC specification occurs in the posterior epiblast by E16 and provide the first insights into the molecular profile of bovine PGCs around day 20 of embryonic development. To our knowledge, this is the earliest timepoint of bovine embryogenesis in which PGCs have been immunolocalized, and the first instance that the specification network of the bovine germline was elucidated. Moreover, this study shows the early migratory path of bovine PGCs through the primitive streak, into the allantois, and through the hindgut endoderm. Having a complete transcriptional profile of early migratory PGCs has shed light on several early aspects of bovine germline development, including proliferative and epigenetic status. In addition to providing further context of the evolutionary convergence and divergence of germline establishment, this knowledge will be useful for the establishment of new ARTs for livestock and humans.

## Notes

### Competing Interest Statement

The authors have declared no competing interest.

### Funding

- USDA Multi-State W5171
- USDA-NIFA Pre-Doctoral Fellowship: 2022-11304

## References

1. Ohinata, Y. et al. Blimp1 is a critical determinant of the germ cell lineage in mice. Nature 436, 207–213 (2005).

2. Magnúsdóttir, E. et al. A tripartite transcription factor network regulates primordial germ cell specification in mice. Nat Cell Biol 15, 905–915 (2013).

3. Yamaji, M. et al. Critical function of Prdm14 for the establishment of the germ cell lineage in mice. Nat Genet 40, 1016–22 (2008).

4. Irie, N. et al. SOX17 is a critical specifier of human primordial germ cell fate. Cell 160, 253–268 (2015).

5. Sasaki, K. et al. Robust in vitro induction of human germ cell fate from pluripotent stem cells. Cell Stem Cell 17, 178–194 (2015).

6. Sasaki, K. et al. The germ cell fate of cynomolgus monkeys is specified in the nascent amnion. Dev Cell 39, 169–185 (2016).

7. Kobayashi, T. et al. Principles of early human development and germ cell program from conserved model systems. Nature 546, 416–420 (2017).

8. Kobayashi, T. et al. Tracing the emergence of primordial germ cells from bilaminar disc rabbit embryos and pluripotent stem cells. Cell Rep 37, 109812 (2021).

9. Sugawa, F. et al. Human primordial germ cell commitment in vitro associates with a unique PRDM14 expression profile. EMBO J 34, 1009–1024 (2015).

10. Yabuta, Y., Kurimoto, K., Ohinata, Y., Seki, Y. & Saitou, M. Gene expression dynamics during germline specification in mice identified by quantitative single-cell gene expression profiling. Biol Reprod 75, 705–716 (2006).

11. Seki, Y. et al. Cellular dynamics associated with the genome-wide epigenetic reprogramming in migrating primordial germ cells in mice. Development 134, 2627–2638 (2007).

12. Planells, B. et al. Gene expression profiles of bovine genital ridges during sex determination and early differentiation of the gonads. Biol Reprod 102, 38–52 (2020).

13. Wrobel, K. H. & Süß, F. Identification and temporospatial distribution of bovine primordial germ cells prior to gonadal sexual differentiation. Anat Embryol (Berl) 197, 451–467 (1998).

14. Soto, D. A. & Ross, P. J. Similarities between bovine and human germline development revealed by single-cell RNA sequencing. Reproduction 161, 239– 253 (2021).

15. Alberio, R., Kobayashi, T. & Surani, M. A. Conserved features of non-primate bilaminar disc embryos and the germline. Stem Cell Reports 16, 1078–1092 (2021).

16. Bergmann, S. et al. Spatial profiling of early primate gastrulation in utero. Nature 609, 136–143 (2022).

17. Castillo-Venzor, A. et al. Origin and segregation of the human germline. Life Sci Alliance 6, e202201706 (2023).

18. Goszczynski, D. E., Denicol, A. C. & Ross, P. J. Gametes from stem cells: Status and applications in animal reproduction. Reproduction in Domestic Animals 54, 22–31 (2019).

19. Goszczynski, D. E. et al. In vitro breeding: Application of embryonic stem cells to animal production. Biol Reprod 100, 885–895 (2019).

20. Botigelli, R. C., Guiltinan, C., Arcanjo, R. B. & Denicol, A. C. In vitro gametogenesis from embryonic stem cells in livestock species: Recent advances, opportunities, and challenges to overcome. J Anim Sci 101, (2023).

21. Mueller, M. L., et al. Germline ablation achieved via CRISPR/Cas9 targeting of NANOS3 in bovine zygotes. Front Genome Ed 5, (2023).

22. Robertson, I., Nelson R., Stringfellow, D. A. & Seidel, S. M. Certification and identification of the embryo. in *Manual of the International Embryo Transfer Society* 103–134 (International Embryo Transfer Society, Savory, IL, 1998).

23. Berg, D. K., van Leeuwen, J., Beaumont, S., Berg, M. & Pfeffer, P. L. Embryo loss in cattle between days 7 and 16 of pregnancy. Theriogenology 73, 250–260 (2010).

24. van Leeuwen, J., Berg, D. K. & Pfeffer, P. L. Morphological and gene expression changes in cattle embryos from hatched blastocyst to early gastrulation stages after transfer of in vitro produced embryos. PLoS One 10, e0129787 (2015).

25. Soto, D. A. et al. Simplification of culture conditions and feeder-free expansion of bovine embryonic stem cells. Sci Rep 11, 11045 (2021).

26. Gokulakrishnan, P., Kumar, R. R., Sharma, B. D., Mendiratta, S. K. & Sharma, D. Sex determination of cattle meat by polymerase chain reaction amplification of the DEAD Box Protein (DDX3X/DDX3Y) gene. Asian-Australas J Anim Sci 25, 733– 737 (2012).

27. Tyser, R. C. V. et al. Single-cell transcriptomic characterization of a gastrulating human embryo. Nature 600, 285–289 (2021).

28. Anderson, R., Copeland, T. K., Schöler, H., Heasman, J. & Wylie, C. The onset of germ cell migration in the mouse embryo. Mech Dev 91, 61–68 (2000).

29. Chen, D. et al. Human primordial germ cells are specified from lineage-primed progenitors. Cell Rep 29, 4568–4582.e5 (2019).

30. Nakamura, T. et al. A developmental coordinate of pluripotency among mice, monkeys and humans. Nature 537, 57–62 (2016).

31. Kojima, Y. et al. GATA transcription factors, SOX17 and TFAP2C, drive the human germ-cell specification program. Life Sci Alliance 4, e202000974 (2021).

32. Di Carlo, A. & De Felici, M. A role for E-cadherin in mouse primordial germ cell development. Dev Biol 226, 209–219 (2000).

33. Clark, J. M. & Eddy, E. M. Fine structural observations on the origin and associations of primordial germ cells of the mouse. Dev Biol 47, 136–155 (1975).

34. Blaser, H. et al. Transition from non-motile behaviour to directed migration during early PGC development in zebrafish. J Cell Sci 118, 4027–4038 (2005).

35. García-Castro, M. I., Anderson, R., Heasman, J. & Wylie, C. Interactions between germ cells and extracellular matrix glycoproteins during migration and gonad assembly in the mouse embryo. J Cell Biol 138, 471–480 (1997).

36. Vejlsted, M., Offenberg, H., Thorup, F. & Maddox-Hyttel, P. Confinement and clearance of OCT4 in the porcine embryo at stereomicroscopically defined stages around gastrulation. Mol Reprod Dev 73, 709–718 (2006).

37. Tam, P. P. & Snow, M. H. Proliferation and migration of primordial germ cells during compensatory growth in mouse embryos. J Embryol Exp Morphol 64, 133– 47 (1981).

38. Donovan, P., Stott, D., Cairns, L., Heasman, J. & Wylie, C. Migratory and postmigratory mouse primordial germ cells behave differently in culture. Cell 44, 831–838 (1986).

39. Cantú, A. V., Altshuler-Keylin, S. & Laird, D. J. Discrete somatic niches coordinate proliferation and migration of primordial germ cells via Wnt signaling. Journal of Cell Biology 214, 215–229 (2016).

40. Grimaldi, C. & Raz, E. Germ cell migration—Evolutionary issues and current understanding. Semin Cell Dev Biol 100, 152–159 (2020).

41. Zhu, Q. et al. Specification and epigenomic resetting of the pig germline exhibit conservation with the human lineage. Cell Rep 34, 108735 (2021).

42. Molyneaux, K. A. et al. The chemokine SDF1/CXCL12 and its receptor CXCR4 regulate mouse germ cell migration and survival. Development 130, 4279–4286 (2003).

43. Hajkova, P. et al. Chromatin dynamics during epigenetic reprogramming in the mouse germ line. Nature 452, 877–81 (2008).

44. Eguizabal, C. et al. Characterization of the epigenetic changes during human gonadal primordial germ cells reprogramming. Stem Cells 2418–2428 (2016).

45. Hill, P. W. S. et al. Epigenetic reprogramming enables the transition from primordial germ cell to gonocyte. Nature 555, 392–396 (2018).

46. Sugimoto, M. & Abe, K. X chromosome reactivation initiates in nascent primordial germ cells in mice. PLoS Genet 3, e116 (2007).

47. Ehrmann, I. et al. An ancient germ cell-specific RNA-binding protein protects the germline from cryptic splice site poisoning. Elife 8, (2019).

48. Elliott, D. J. et al. An evolutionarily conserved germ cell-specific hnRNP is encoded by a retrotransposed gene. Hum Mol Genet 9, 2117–2124 (2000).

49. Ghieh, F. et al. Cryptic splice site poisoning and meiotic arrest caused by a homozygous frameshift mutation in RBMXL2: A case report. Andrologia 54, (2022).

50. Siachisumo, C., et al. An anciently diverged family of RNA binding proteins maintain correct splicing of a class of ultra-long exons through cryptic splice site repression. bioRxiv 2023.08.15.553384 (2024) doi:10.1101/2023.08.15.553384.

51. Hikabe, O. et al. Reconstitution in vitro of the entire cycle of the mouse female germ line. Nature 539, 299–303 (2016).

52. Zhou, Q. et al. Complete meiosis from embryonic stem cell-derived germ cells in vitro. Cell Stem Cell 18, 330–340 (2016).

53. Sakai, Y. et al. Induction of the germ cell fate from pluripotent stem cells in cynomolgus monkeys. Biol Reprod 102, 620–638 (2020).

54. Gkountela, S. et al. The ontogeny of cKIT+ human primordial germ cells proves to be a resource for human germ line reprogramming, imprint erasure and in vitro differentiation. Nat Cell Biol 15, 113–22 (2013).

55. Guo, F. et al. The transcriptome and DNA methylome landscapes of human primordial germ cells. Cell 161, 1437–52 (2015).

56. Gomes Fernandes, M., Bialecka, M., Salvatori, D. C. F. & Chuva de Sousa Lopes, S. M. Characterization of migratory primordial germ cells in the aorta-gonad-mesonephros of a 4.5-week-old human embryo: a toolbox to evaluate in vitro early gametogenesis. Mol Hum Reprod 24, 233–243 (2018).

57. Shirasawa, A. et al. Efficient derivation of embryonic stem cells and primordial germ cell-like cells in cattle. Journal of Reproduction and Development **Epub**, 2023–087 (2024).

58. Mitsunaga, S. et al. Relevance of iPSC-derived human PGC-like cells at the surface of embryoid bodies to prechemotaxis migrating PGCs. Proc Natl Acad Sci U S A 114, (2017).

59. Vijayakumar, S. et al. Monolayer platform to generate and purify primordial germ-like cells in vitro provides insights into human germline specification. Nat Commun 14, 5690 (2023).

60. Chitiashvili, T., Hsu, F., Dror, I., Plath, K. & Clark, A. FGFR3 is expressed by human primordial germ cells and is repressed after meiotic initiation to form primordial oocytes. Stem Cell Reports 17, 1268–1278 (2022).

61. Chuva de Sousa Lopes, S. M., Roelen, B. A. J., Lawson, K. A. & Zwijsen, A. The development of the amnion in mice and other amniotes. Philos Trans R Soc Lond B Biol Sci 377, (2022).

